# Ensembling Unets, sparse representation and low dimensional visualization for rare chromosomal aberration detection in light microscopy images

**DOI:** 10.1101/2023.09.11.557124

**Authors:** Antonin Deschemps, Eric Grégoire, Juan S. Martinez, Aurélie Vaurijoux, Pascale Fernandez, Delphine Dugue, Laure Bobyk, Marco Valente, Gaëtan Gruel, Emmanuel Moebel, Mohamed Amine Benadjaoud, Charles Kervrann

## Abstract

In biological dosimetry, a radiation dose is estimated using the average number of chromosomal aberration per peripheral blood lymphocytes. To achieve an adequate precision in the estimation of this average, hundreds of cells must be analyzed in 2D microscopy images. Currently, this analysis is performed manually, as conventional computer vision techniques struggle with the wide variety of shapes showcased by chromosomes. The false discovery rate of current automated detection systems is high and variable, depending on small variations in data quality (chromosome spread, illumination variations …), which makes using it in a fully automated fashion impossible. Automating chromosomal aberration is needed to reduce diagnosis time. Furthermore, an automated system can process more images, which improves confidence intervals around the estimated radiation dose. We build an object detection model to automate chromosomal aberration detection using recent advances in deep convolutional neural networks and statistical learning. We formulated the problem of rare aberration detection as a heatmap regression problem requiring the minimization of a sparsity-promoting loss to reduce the false alarm rate. Our Unet-based approach is analoguous to a one-stage object detector, and keeps the number of hyperparameters to a minimum. Finally, we demonstrate large performance improvements using an ensemble of checkpoints collected during a single run of training. A PCA-based strategy is used to provide cues for interpretation of our deep neural network-based model. The methodology is demonstrated on real, large, and challenging datasets depicting rare chromosomal aberrations and is favorably compared to a reference dosimetry technique.

## 1 Introduction

Because ionizing radiation causes chromosomal aberration in peripheral blood lymphocytes, the average number of aberration per lymphocyte can be used to estimate a radiation dose after an exposure event. This is especially relevant for accidental (or potentially criminal) exposures, where readings from a dosimeter may not be available. According to the IAEA, the current gold standard for biological dosimetry is the counting of dicentric chromosomes [1], i.e., chromosomes with two centromeres. To observe those aberrations, blood cells are grown for 48 hours, and Demelcocine is used to stop cell division in the metaphase step of mitosis [3]. Because of this, an image of a single cell undergoing metaphase (where the 23 pairs of chromosomes are visible) is often called a metaphase. As chromosome are translucent objects, Giemsa staining is used to increase contrast as the sample is spread on a glass slide for imaging. Figure 1 provides a chronological overview of the image acquisition process. A calibration curve links the aberration rate with an ionizing radiation dose, as displayed in Figure 1c). The rate of aberration for an individual who was not exposed to any radiation is around 1 aberration every 1000 cells. Therefore, controlling the false discovery rate (FDR) of aberrations is essential, to avoid over-diagnosing acute radiation exposition and overloading care facilities, especially in the case of a large scale exposition.

**Figure 1.**
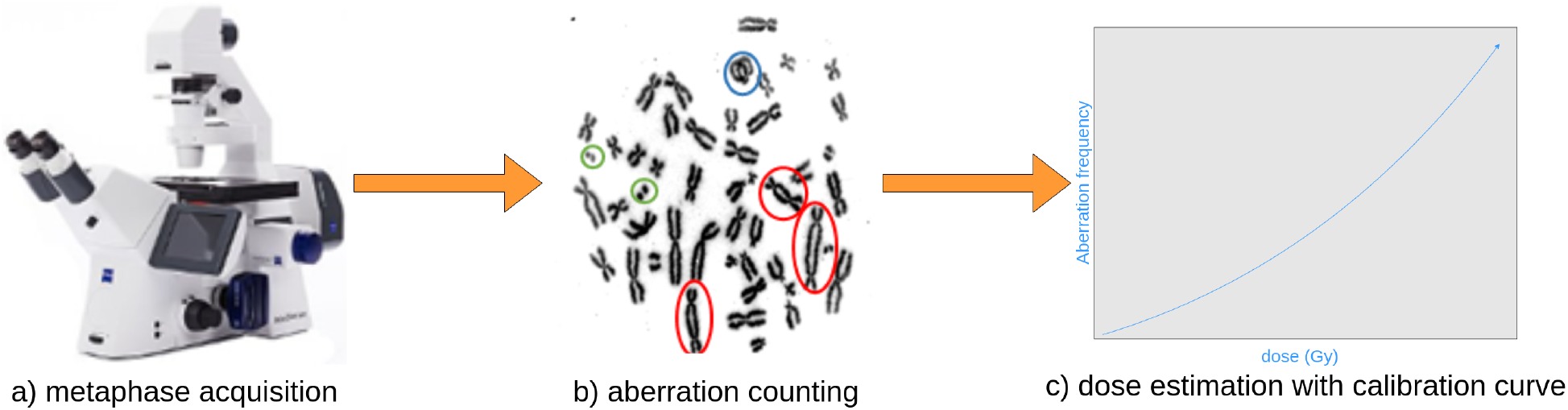
Steps required to estimate a radiation dose in biological dosimetry: a) (automated) metaphase acquisition; b) dicentric chromosomes (red circle), and fragments (greeen circle) counting; c) dose regression from aberration counts.

While metaphase acquisition can be partially automated with current tools, dicentric chromosome counting still relies on human expertise, mainly because a monocentric chromosome can be deformed in such a way that it looks like a dicentric chromosome. Disambiguating between monocentric and dicentric chromosomes may be achieved by shifting the focus plane of the microscope, as monocentric chromosomes with overlapping telomeres can be discriminated from dicentric chromosomes based on variations in thickness (see Figure 1b) for an example of dicentric chromosome, circled in red). Unfortunately the images acquired by the microscope camera are 2D which makes this thickness information unavailable.

### 1.1 Supervised learning and regression model ensemble for aberration counting

Aberration counting is routinely performed on 2D images but still remains a tedious task, even it assisted by semi-automatic methods. To improve on the performance of current automated dicentric counting methods, we investigated recent advanced methods in deep learning. Since the seminal ImageNet paper [18], convolutional neural networks (ConvNets) trained with stochastic gradient descent (SGD) have brought significant advances to many common tasks in computer vision, like image classification, object detection or image segmentation. Modern ConvNets are easy to train thanks to batch normalization and skip connections. However, deep learning models have been shown to suffer from over-confidence and mis-calibration. In other words, if a classification model predicts a class with a probability of 99%, there is no guarantee that the error rate is close to 1%, see [13].

To overcome this difficulty, we have investigated the human decision making process. For any specific slide, if one asks several trained human experts to count chromosomal aberrations, the agreement between experts is not perfect, even with the additional 3D shape information obtained by shifting the focus plane of the microscope. In case of disagreement, experts discuss their disagreement and make a consensus decision. We can emulate this strategy using a set (also called “ensemble”) of models providing different predictions. Each artificial expert (i.e., model) “votes” for any given object observed in the metaphase image. The model votes for a region of the image if its output exceeds a pre-specified confidence threshold. Objects getting a minimal number of votes are considered to be plausible aberrations. No architectural modification is needed, and significant performance improvements are achieved. During model training, SGD samples the parameter space of the model. SGD samples are retrieved at the end of each epochs, and the ensemble is built from a random sample of those checkpoints retrieved during training.

Because we exploit randomness to build a diverse ensemble, it is helpful to visualize training dynamics to better interpret the relationships between training stochasticity and ensemble diversity. Moreover, visualizing training dynamics of deep neural networks is known to be difficult, as this class of model is usually over-parameterized. A common solution is to display the output of a network instead of its parameters. For tasks like image classification, this output is a single vector which makes common dimensionality reduction techniques very effective so that one can easily visualize class separation across training epochs. For dense tasks, a prediction is made for every single pixel of the input image instead. Therefore, we consider this feature map as a set of independent vectors to emulate common approaches in training visualization for classification models. This makes visualization of class separation during training possible, and provides a visual explanation for the performance gains brought by our aggregation procedure.

### 1.2 Contributions

In this paper, we propose an aggregation-based method for rare chromosomal aberration detection. Aggregation solves two key issues. First, it allows one to build a high-performing chromosomal aberration detector out of several instances of a simple Unet model, without the need for extensive tweaks or architectural modifications. Second, it reduces uncertainty around model performance. During the training of a single model, the stochasticity of SGD will lead to variations in test performance. We show that the variation in test performance between ensembles of randomly sampled checkpoints is low. In fact, over a set of randomly sampled ensembles, it is lower than between single checkpoints collected during training. Finally, we show promising results on calibration curve estimation.

### 1.3 Organization

The remainder of the paper is organized as follows. In Section 2, we first describe the related works for automated biological dosimetry. Second, we briefly summarize the main features of ConvNets and review the ensemble approaches in deep learning. In Section 3, we present our training framework based on sparse representations and model ensemble. In this section, we describe a novel visualization approach to explore the model building during training. Section 4 is devoted to data descriptions and evaluation metrics definition. We use two datasets: a training dataset where aberrations are overrepresented, and a validation dataset for calibration curve estimation. In Section 5, we evaluate the performance the single model and model ensemble and demonstrate the robustness of the method regarding the presence of debris in metaphase images. Furthermore, we examine the performance our model given the distribution shift between our imbalanced training dataset and a more representative testing dataset. Simple modifications on thresholding can be used to handle this shift and estimate a realistic calibration curve in spite of the unbalanced training dataset. In Section 6, we sum up our results and discuss perspectives and future work in biological dosimetry.

## 2 Related works

In this section, we present the related works that served as a starting points for developing our method.

### 2.1 Cytogenetic biological dosimetry

As chromosomes are a very common object of interest in cytogenetic biological dosimetry, characterizing the shape of chromosomes with computer vision tools has been a longstanding area of research. Typically, one approach consists in detecting chromosome centromeres [39], as they are a reliable indicator to separate dicentrics from monocentrics. Centromeres may be retrieved by locating minimas of the chromosome width along the centerline, as shown in [40, 37].

Those classifiers have been used in pipelines tackling automated dose estimation in several different commercial solutions. In Europe, Metafer (provided by MetaSystems) has the largest marketshare, and its performance have been evaluated several years ago [41, 10]. In [24], Liu et al. showcased a software stack called ADCI which estimated a dose from metaphase images, using conventional computer vision techniques and prior chromosome morphology. ADCI uses a significant degree of prior knowledge to localize and classify dicentric chromosomes. For example, various degrees of chromosome condensation are identified based on cell entry into metaphase, and late metaphases are rejected to avoid chromosomes with excessively separated telomeres, which leads to spurious detections of dicentric chromosomes. Chromosomes are segmented, and metaphases are accepted or rejected based on the number of objects they contain [33]. Centromeres are identified to discriminate dicentrics from monocentrics. While ADCI reaches a high level of performance, it deals with the variations in image quality and chromosome morphology by using sophisticated image selection models.

More recently, supervised deep learning has been used to tackle the DC detection problem. In [16], the authors demonstrate that Faster R-CNN can be used to detect DCs in metaphases. This model is thoroughly evaluated in [17].

### 2.2 Key-point regression in deep learning

In key-point regression, Gaussian spots are predicted over specific landmarks of the image, like the eyes on a human face or the joints of a skeleton. It is a subtask of a large number of computer vision tasks like facial recognition or pose estimation [4, 5]. It can be solved with common image segmentation models by minimizing the *L*_2_ distance between a predicted heatmap and the corresponding ground truth over a training dataset. While heatmap regression and object detection are different tasks at first glance, and heatmap regression does not predict bounding box extent, several authors have proposed a unified method that consists in predicting a center point and bounding box dimensions [44], or the corners of a bounding box as key-points [21].

### 2.3 Object detection and counting

Object counting is a common computer vision task, and a wide variety of solutions have been suggested in the literature. Density-based methods aim to predict counts by integrating a density map [22, 7]. This class of methods is usually simple to implement and reaches a high level of performance. However, it does not fit bounding boxes, and summary statistics like Precision and Recall cannot be computed. Alternatively, counting can be solved as a subtask of object detection, by enumerating the bounding boxes belonging to a certain class. However, accurately predicting bounding boxes is more difficult than predicting a density map, and detection-based methods usually perform worse than density-based ones [6]. In [20], the authors propose a detection-based method that relies on predicting a Gaussian spot centered on the object, but does not predict bounding box extent. In [21], the author propose a model that uses the same heatmap-based technique, but also predicts a (height, width) tuple for a bounding box.

### 2.4 Ensemble and approximate Bayesian deep learning

#### 2.4.1 Model aggregation

A large number of papers have studied ensemble methods for neural networks, either to improve model calibration, for uncertainty modeling or to improve performance. The authors focus either on *sampling* a diverse set of models to build an ensemble, or on the properties of the aggregated prediction.

For classification models, Lakshminarayanan et al. [19] showed that ensembles of neural networks improved on the performance and calibration of single models. As an ensemble of *M* models requires *M* training runs, [14] showed that a carefully chosen learning rate schedule could encourage loss landscape exploration to get a collection of models with high diversity in a single training run by retrieving checkpoints. In [42], samples and checkpoints are re-weighted according to their performance, like AdaBoost [35].

In semantic segmentation, ensemble methods have received attention because they improve performance and provide a localized measure of uncertainty, usually by computing the entropy of the ensemble average for every pixel in the image. In [27], an ensemble of fully convolutional neural networks is used to segment aerial images. In [36], a diverse ensemble is built by training several Unets [34] with different encoders, and predictions are aggregated with a weighted average. In [29], the authors provide a review of ensemble methods for polyp segmentation.

For bounding-box based detection models, aggregation is required, as several boxes localizing the same object may overlap. Non-Maximum Suppression (NMS) [30] is then used and consists in sorting the boxes with respect to their confidence levels. Lower confidence boxes overlapping a high confidence box beyond a specific IoU (Jaccard index) threshold are discarded. In [38], box merging algorithms are explicitly considered in a particular model aggregation framework. The authors suggest computing an “average” box by weighting coordinates based on the box confidence.

#### 2.4.2 Approximate Bayesian deep learning

Common techniques used to sample the posterior distribution are intractable for modern neural networks given their large parameter counts. In [26], Mandt & al demonstrate that SGD can be seen as an Orstein-Uhlenbeck process with some limit Gaussian distribution. SGD can be seen as Langevin sampling of the posterior weight distribution, and SGD samples can be used to estimate the mean and covariance of the Gaussian posterior distribution. This provides a cheap and simple way to retrieve samples of this distribution during the training procedure.

in [11], Garipov & al show that local minima are connected by low-loss paths, and that one could average weight vectors along deterministic trajectories between those local minima to improve performance. In [15], Izmailov & al confirm the theoretical analysis of [26] by showing that averaging SGD iterates leads to wider minima and improved generalization in practice.

Using the theoretical insights explained in [26], Maddox et al. [25] approximate the limit posterior weight distribution with a Gaussian distribution, where the covariance matrix is defined as the covariance of the last gradient descent iterates. New sets of weights can be sampled from this posterior distribution for ensemble and uncertainty estimation. In [43], the authors go further by sampling an ensemble from several modes of the posterior weight distribution to increase ensemble diversity and therefore performance.

Finally, other approximations of the posterior weight distribution have been proposed. In [9], the authors use Kalman filtering to derive a sequential estimate of the posterior distribution over the weights.

## 3 Methods

In this section, we first give a precise definition of the data and model, including the loss for training. Second, we explain our aggregation procedure, going from a set of continuous heatmap predictions for a single image to a set of binary decisions maps and finally, to an aggregated ensemble-level prediction. Finally, we explain the PCA-based visualization techniques used to justify the performance gains of the ensemble and its robustness to the presence of debris in metaphase images.

### 3.1 Keypoint regression with heatmap regression models

In Heatmap Regression Models (HRMs), objects of interest are represented as Gaussian spots. The model is trained to predict spot positions in the image domain, with a labelled dataset 𝒟 = {(*x*_1_, *y*_1_), *· · ·*, (*x*_*n*_, *y*_*n*_)} comprised of *n* realizations of a pair of random variables (*X, Y*). For each image *x*_*i*_, we have *x*_*i*_(*u, v*) ∈ [0, 1] at each location (*u, v*) ∈ Ω, where Ω denotes the image grid of size |Ω| = *H × W*.

#### 3.1.1 Model design and sparsity promoting loss function

Our heatmap regression model is a convolutional neural network *ϕ*_*θ*_(*x*) : *x* ∈ [0, 1]^*H×W*^ *→ y* ∈ [0, 1]^*H×W*^. The final output is constrained between 0 and 1 with a sigmoid layer. We use the Unet architecture [34] to predict a low resolution heatmap. The image *y*_*L*_ is of size (*H/L, W/L*) for some arbitrary downsampling factor *L*, as the location accuracy provided by the highest resolution output is not useful. For Unet-based architectures, images are usually downsampled (or upsampled, in the decoder) by a factor of 2 at each layer: at layer *l* of the encoder, features have a spatial size of *H/*2^*l*^ *× W/*2^*l*^. For the sake of simplicity, notations are given in the single-channel case. Additional classes of aberrations (like fragments) are modeled with additional channels, so that a third index is added. In the remainder of the paper, we consider two aberration classes, dicentric chromosomes and fragments. The parameters *θ* are learned by solving the following optimization problem:

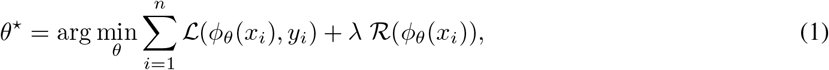

where *λ* is an hyperparameter that balances the data fidelity term and the regularization term. In our modeling approach, the data fidelity term *ℒ* has the following form:

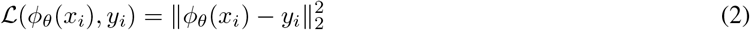

Because the number of aberrations is very low compared to the number of pixels in the image, the background is expected to be 0, except in a small number of “hot” spots corresponding to locations containing aberrations. The Sparse Variation (SV) regularizer (3) has been specifically considered here to encourage the emergence of a very small number of “hot” spots as aberrations are rare events in GIEMSA images. This regularizer is defined as [32]:

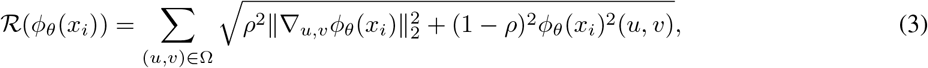

where *ρ* is a parameter that balances the sparsity and the smoothness terms in the predicted heatmap. The components of the gradient vector are computed with respect to the image coordinate axes as follows:

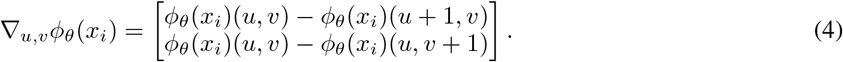

As this is a simple linear transformation of the image, computational overhead is minimal.

The criterion (2) is highly non-convex because of non-linearities in *ϕ*_*θ*_. Therefore, finding a global minimum is hopeless. Nevertheless, a good local minima may be found using iterative first order methods, usually some variant of SGD. The exact gradient of the training criterion with respect to *θ* is estimated on a random subset *𝒥* of the complete dataset *𝒟*, because of memory constraints:

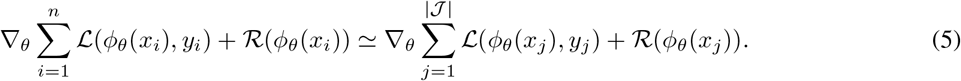

In Figure 3, we reported two curves corresponding the cumulative distributions functions of the total variation images computed over the prediction maps regularized with the sparse variation regularizer (blue curve) and without (red curve). See 2 for an example of images on which this total variation is computed.

**Figure 2.**
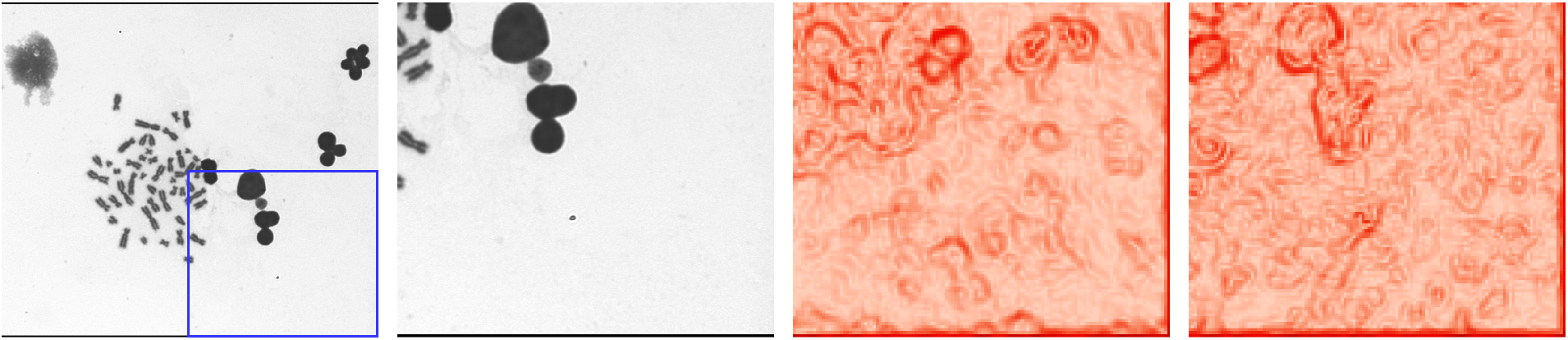
Visual comparison between regularized and unregularized model. First image from the right: input image, second: bottom right crop, third: gradient norm of the prediction for the unregularized model, fourth: gradient norm of the prediction for the regularized model.

**Figure 3.**
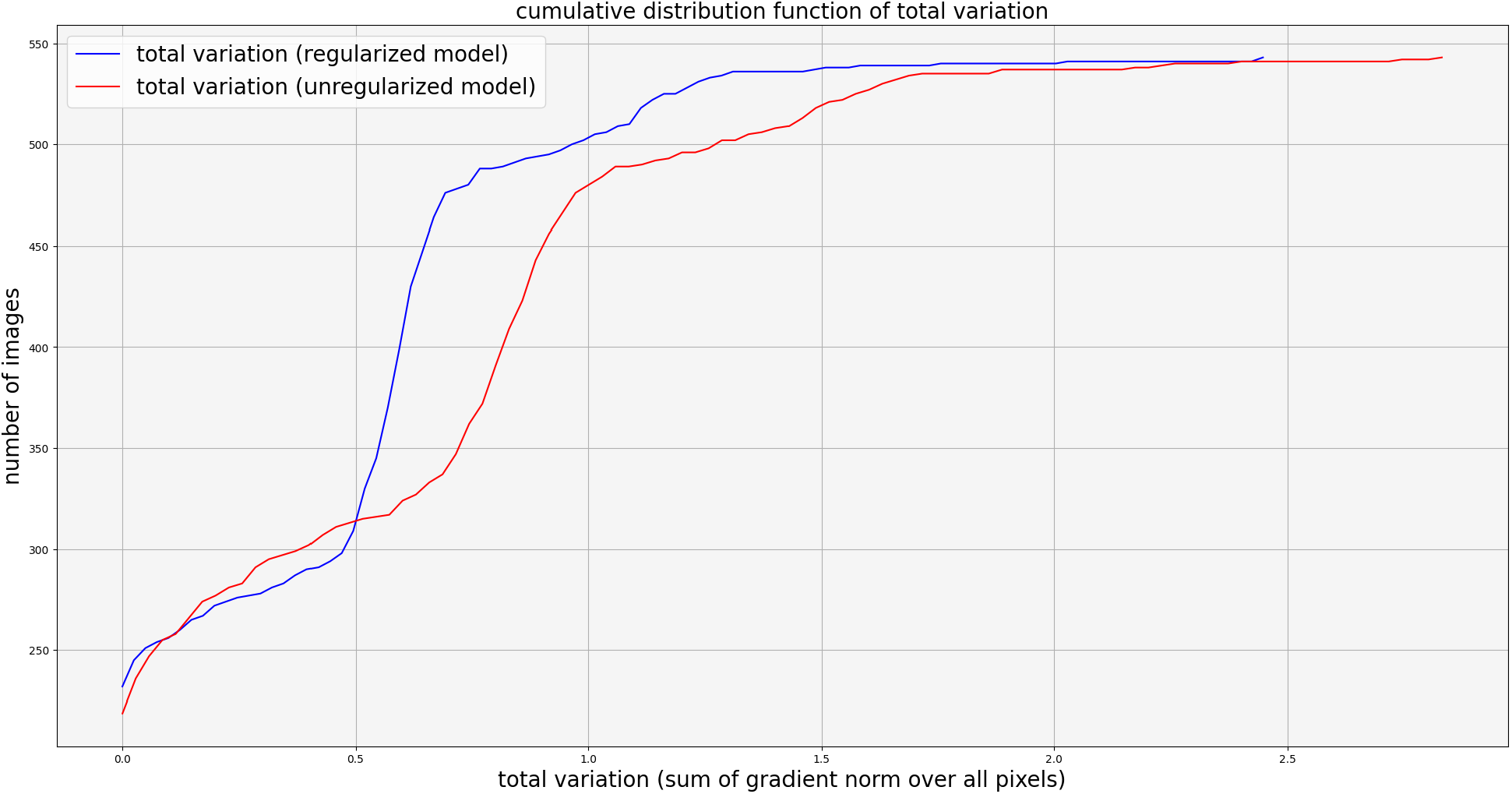
For each image, we compute the total variation of its prediction for the regularized and unregularized model, as shown in Figure 2. This figure represents the Cumulative Distribution Function (cdf) of the total variation over all images in the test set.

### 3.2 Implicit ensembling of neural networks

To reduce the number of false positives (i.e, improve Precision) at a fixed recall level, we investigated the ensemble method introduced in [19]. Because training a deep neural network is a stochastic process, successive training runs of the same model tend to explore different regions of the parameter space. Those local minimas are usually very close in terms of validation loss, but their predictions are not identical. This behavior has received significant attention in the literature [14, 11]. As shown in [14], it may not even be necessary to run several successive training runs. A carefully chosen learning rate schedule may be enough to achieve enough parameter space exploration to build a diverse ensemble from checkpoints of a single training run. In our case, we even find that the gradient noise introduced by stochastic batch sampling leads to sufficient checkpoint diversity for agregation to be worthwile without any specific learning rate schedule. Therefore, we do not have to deal with the training instability mentioned in [25].

More formally, SGD can be seen as a Langevin sampling of the posterior weight distribution *p*(*θ*| *𝒟*) [26]. To avoid excessive autocorrelation between samples, we store the vectors *θ* at the end of each epoch, instead of every gradient step. This can be interpreted as a variant of *thinning*, also used in Monte Carlo Markov Chain inference. For each image *x*_*i*_, *i* ∈ {1, *· · ·, n*} in the test set, we consider a set 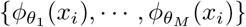 of predictions, where *M* is the number of predictions (or models). Using a confidence threshold *T*_*C*_, we build a set 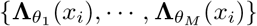 of *M* different binary predictions for each test image *x*_*i*_, that we sum over all members of the ensemble at each location (*u, v*) ∈ Ω as follows:

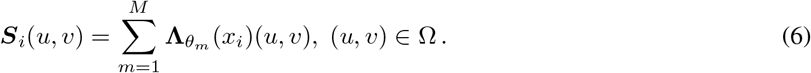

Finally, we set an agreement threshold *T*_*A*_ to compute the agreement between the artificial “experts”. The setting of thresholds *T*_*A*_ and *T*_*C*_ impact the final decision. If the confidence threshold *T*_*C*_ is high and the voting threshold *T*_*A*_ is low, the decision will be made from a small set of experts. Otherwise, a small value of *T*_*C*_ but a high voting threshold *T*_*A*_ means that low confidence predictions are considered, but a higher agreement between them is needed to confirm a detection. In the end, we get a precision surface depending on *T*_*C*_ and *T*_*A*_. Our aggregated decision for any image *x*_*i*_ is a binary image ***D***_*i*_ such that value at location (*u, v*) is 0 if no aberration is predicted, and 1 otherwise (See Figure 4 for illustration):

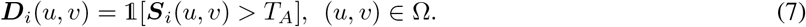

**Figure 4.**
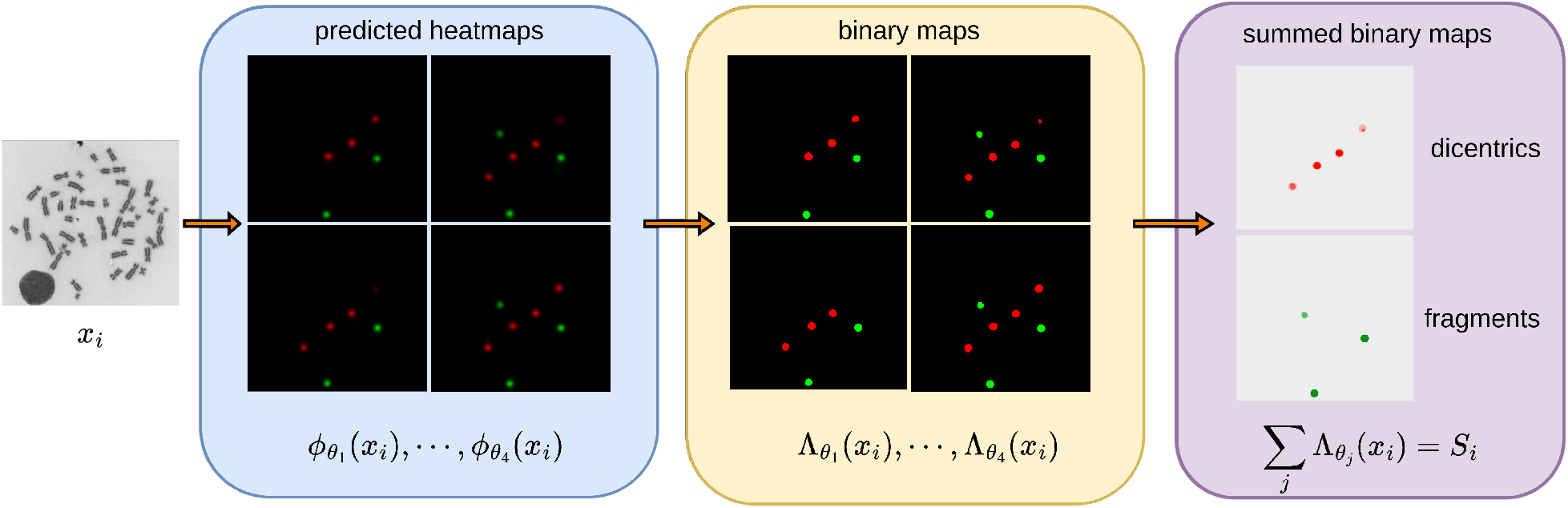
Vote-based aggregation of checkpoints. Dicentric chromosomes are plotted in red, and fragment predictions are plotted in green. For every image *x*_*i*_, the heatmap prediction *ϕ*_*θ*_(*x*_*i*_) is binarized (giving **Λ**_*θ*_(*x*_*i*_)) with a confidence threshold *T*_*C*_. Those maps are summed (giving *S*_*i*_), and regions of the image receiving more than *T*_*A*_ votes are considered as detections. Darker shades of red and green indicates region of the images receiving more votes.

The agreement threshold *T*_*A*_ can be adjusted by the end user to optimize either Precision or Recall scores, like the confidence threshold *T*_*C*_. We discuss the effects of choosing a specific threshold in Section 4.2.2.

### 3.3 Setting of model parameters

We trained Unet for *N*_*e*_ = 100 epochs epochs with Adam [8], with a constant learning rate of 3 *×* 10^*−*4^, a weight decay parameter of 0.1 and a batch size of 12 on a single Tesla V100. The learning rate was unchanged during training to ensure parameter space exploration, using an analoguous reasoning to the one provided in [15].

We did not use data augmentation for two reasons. First, we found that the wide variety of chromosome morphology and orientations in our dataset was enough for our model to learn this invariance. We did not observe detection failures based on object orientation. Second, more agressive data augmentation like noise or blurring quickly made monocentrics and dicentrics indistinguishable. Training stability was very sensitive to the variance of the blurring kernel or the Gaussian noise, because accurate chromosome classification relies on very small details.

We predict a lower resolution heatmap of size *H*^*′*^ = 224, *W*^*′*^ = 252, where height and width are downsampled by a factor of 4. While batch sampling (and therefore parameter space exploration) is randomized, parameter initialization is fixed between training runs. We ran a grid search with log_10_ spacing for *λ* regularization parameter with 10 and 10^*−*4^ as upper and lower bound of the search interval. Training was implemented in PyTorch [31], and uses segmentation_models_pytorch implementation of Unet.

### 3.4 Visualization of the training dynamics of single model

As an additional visual explanation for aggregation performance gain, it may be informative to display training trajectories in feature space. Classification networks output a single classification vector per image, so that plotting training dynamics over time using dimensionality reduction is relatively easy (see [23]). A scatterplot of classification vectors embedded in a lower dimension at each epoch provides a good view of how classes are progressively separated during training. Usually, UMAP [28] is used, which requires a pairwise distance matrix between classification vectors. However, this visualization does not work for models outputting a probability distribution for all locations *u, v* of the input image, like Unet.

To address this issue, we adapt the approach [23] to our context. We consider feature maps as bags of independent feature vectors. Figure 5 provides a visual summary of our approach. We do not retain feature vectors for all locations *u, v* in the feature map. Instead, we only select feature vectors corresponding to the locations of aberrations, and retrieve some feature vectors at random ‘background’ (i.e., where there are no aberrations) locations. This bag of features is projected on the 2D plane using its PCA decomposition. By retrieving the same locations across several training steps, we can visualize how the both aberration classes and the background are separated during training. An SVM classifier fitted on the embeddings retrieved for a single epoch is used to map regions of the latent space to a specific class, which helps visualize the dynamics of training. The rest of this section gives a formal overview of our visualization technique.

**Figure 5.**
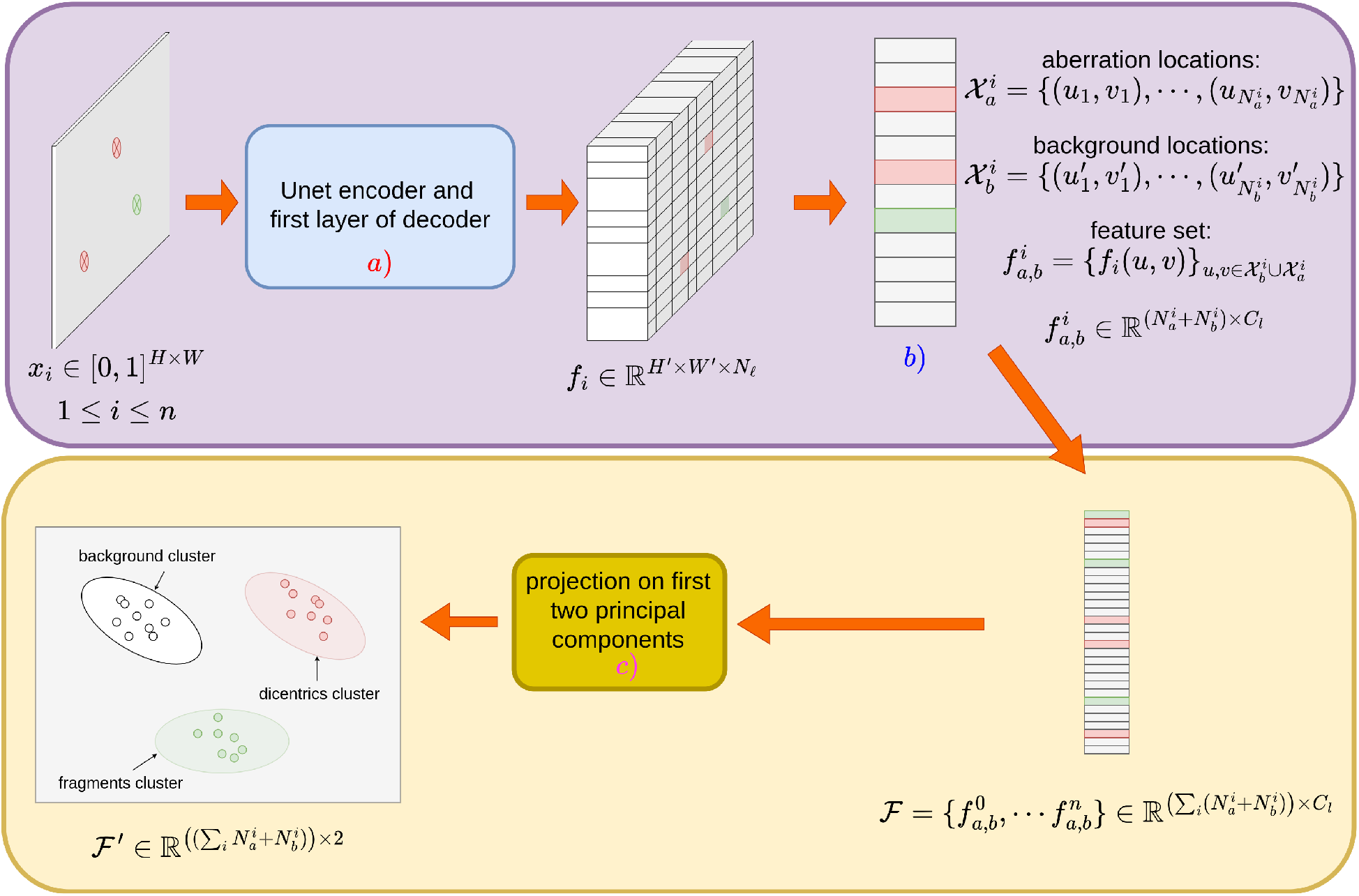
Procedure used to display feature separation in the latent space of the last decoder block for a single epoch (i.e a single weight vector *θ*. For all images *x*_1_, *· · ·, x*_*n*_, the feature maps produced by the last layer of the decoder are retrieved and treated as a set of independent, *C*_*l*_-dimensional feature vectors. Using PCA dimension reduction, we produce a 2D scatterplot that shows how the model separates the different classes (background, dicentrics, fragments) across training epochs. Note that the eigenvectors used for this dimensionality reduction are computed over *all* epochs of training.

Formally, for an input image *x*_*i*_ of size *H × W*, the *ℓ*-th layer of our Unet produces a feature volume 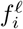 of size *H*^*′*^ *× W*^*′*^ *× N*_*ℓ*_ (see Section 5.3 and Figure 5 (a)). Here, we choose the second-to-last layer of the decoder and we drop the superscript *ℓ* to improve readability, so that 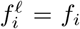 For the last layer, *N*_*ℓ*_ = 128 (see illustration in Figure 5). We retrieve the set of feature vectors that correspond to the spatial locations of aberrations (dicentrics and fragments) in image *x*_*i*_. For a set of 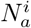 aberrations located at 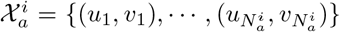 in image *x*_*i*_, we build a set of feature vectors 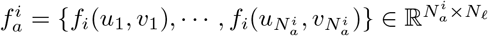. We retrieve the feature vectors corresponding to all aberrations in every image of the test set. We also sample an additional set of 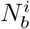background pixels, denoted as 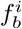 at locations 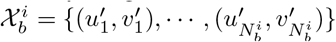 Those locations are randomly sampled, provided the locations do not correspond to aberration pixels. Therefore, they correspond to background, monocentric or debris. Finally, we define 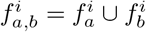 so that the total number of feature vectors in 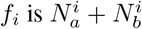, as shown in Figure 5b). It is worth noting that 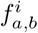 is a subset of the complete feature map *f*_*i*_. This makes the visualizations described less cluttered, and reduces computation time. Finally, this feature set 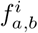 is retrieved for each image *x*_*i*_ in the test set, to build a large feature set 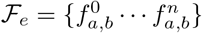 where *e* corresponds to the set of model parameters retrieved at epoch *e*.

The set of feature vector *ℱ*_*e*_ is retrieved for each epoch *e ∈* {1, *· · ·, N*_*e*_}. These sets are concatenated in global set 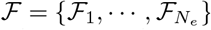 Although all the subsets of *ℱ* correspond to the same locations, they are different at each epoch *e* because of the stochasticity of gradient descent. PCA can be used to visualize those feature sets as 2D scatterplots, and in particular to check how well aberrations are separated from the background at every epoch. The set *ℱ*_1_ is chosen as a reference feature set and *ℱ*_*e*_, *e* ∈ {2, *· · ·, N*_*e*_} is registered with respect to *ℱ*_1_ using the Procrustes method [12]. This guarantees that latent space scale shifts or rotations are removed for visualization. A PCA decomposition is computed on *ℱ*, and for each epoch *e*, each feature set in *ℱ*_*e*_ is projected on the first two principal components of this decomposition. This provides a 2D visualization of the trajectory of each feature vectors during training.

Furthermore, we train a kernel SVM classifier *p*_*e*_ on the 2D embeddings of *ℱ*_*e*_ to predict which aberration class corresponds to a location in 2D embedding space. This classifier takes a 2D embedding (a vector of *ℱ*_*e*_) as an input, and outputs a probability distribution over three classes: background, dicentric chromosome and fragment. For all epochs 1 *≤ e ≤ N*_*e*_, we train a different classifier, and predict a probability distribution over a grid that samples the 2D aberration space uniformly. For all positions (*u, v*) of this grid, a probability distribution over the total number *N*_*r*_ of aberration classes *p*_*e*_(*u, v, r*) ∈ [0, 1], *r* ∈ {0, *· · ·, N*_*r*_},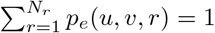 is predicted at epoch *e*. As all the point clouds are aligned and a single set of principal components is computed for all time steps, the changes in the decision boundary from one epoch to the next can be solely attributed to the dynamics of training.

Finally, to visualize the displacement of class boundaries across training, we define the averaged classifier:

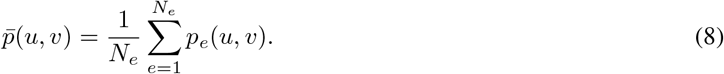

Visualizing the spread of the distribution of 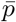 can be done by computing the entropy of the distribution predicted by the averaged classifier:

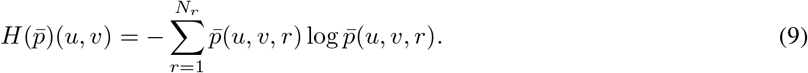

## 4 Materials

### 4.1 Data description

Our training dataset is composed of 5430 labelled images of size *H* = 888, *W* = 1008, padded to *H* = 896, *W* = 1008 to ensure that downscaling has an integer height and width. Labels are binary images with size *H*^*′*^ = 202, *W*^*′*^ = 252, taking value 0 everywhere except at the center of chromosomal aberrations (roughly between the two centromeres for a dicentric chromosome), where it takes value 1. There is one binary image per aberration classes for each image, so that aberration classification is possible. Chromosomal aberrations are the only labelled objects, neither debris nor monocentric chromosomes are labelled. We chose this labelling scheme instead of semantic segmentation or bounding boxes as it lead to the lowest labelling time, which in turn meant a greater number of images could be labelled for the same labelling budget. For the same reason, we did not label debris or monocentric chromosomes. This also prevented the discovery of trivial models where chromosomes would be detected but always labelled as monocentric, as they outnumber dicentric ones by an extremely large margin. As explained in Section 3.1, this binary image is blurred with a Gaussian kernel, to reduce the underrepresentation of the labels against the background.

Images have been selected so that each images contains only the chromosomes corresponding to a single cell, i.e., no image contains more than 46 chromosomes. The metaphases in the dataset do not have any missing chromosome, or an excessive chromosome count. Our dataset contains 5021 dicentrics and 7540 fragments, Figure 6 shows the repartition of images of images into aberration counts bins. Images with a high aberration count are much rarer than images with a low aberration count. On average, there is more than one aberration per image, which corresponds to a very high ionizing radiation dose. As a normal metaphase contains 23 chromosome pairs, this means that even in this case, the overwhelming majority of chromosomes are healthy (i.e monocentric) ones. In our evaluation setting, the training set consists of 80% of those image. 10% of the images are retained for a validation set, used for hyperparameter selection. Finally, 10% of the data is held as a test set for a fair performance evaluation.

**Figure 6.**
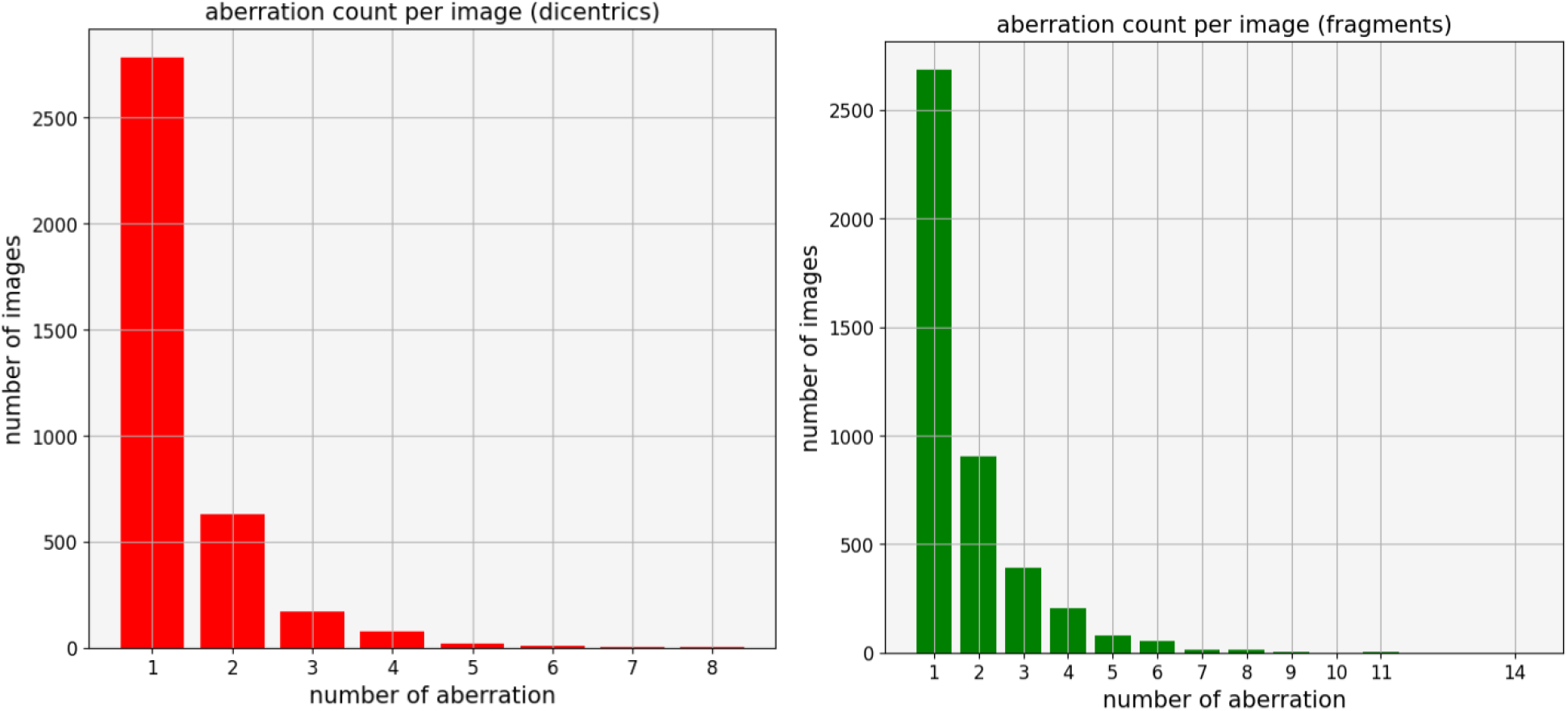
Repartition of images into aberration counts bins.

As this training dataset is not representative of a real-world exposition, we use another dataset to estimate the calibration curve of our model. This dataset was built by collecting metaphases from samples irradiated at specific, known doses. The aberrations in this dataset are not labelled. It contains 21215 metaphases taken from samples irradiated at 0 Gy, 0.1 Gy, 0.2 Gy, 0.3 Gy, 0.5 Gy, 0.7 Gy, 0.9 Gy, 1 Gy, 1.5 Gy, 2 Gy, 3 Gy and 4 Gy.

### 4.2 Evaluation metrics

#### 4.2.1 Performance of a single model

While our predicted heatmap *ϕ*_*θ*_(*x*_*i*_) can take any value between 0 and 1 at each spatial position (*u, v*) in the image, ultimately a binary decision needs to be taken with regard to the presence or absence of aberration at location (*u, v*) ∈ Ω. In the next step *ϕ*_*θ*_(*x*_*i*_) is used to build a binary map **Λ**_*θ*_(*x*_*i*_) given an arbitrary threshold *T*_*C*_:

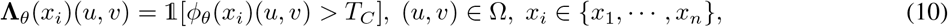

where 1[.] is the indicator function. As *ϕ*_*θ*_ is trained to predict a Gaussian spot, thresholded predictions are binary images comprised of approximately circular spots. Furthermore, the Gaussian spots in the ground truth heatmap are also converted to binary circles with a fixed threshold *T*_*GT*_, which is set to a small fixed value (e.g., 0.01).

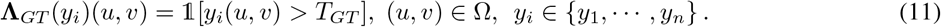

Once a binary decision for the presence of an object is made at each location in the image, we define True Positives, false positives and false negatives. In object detection, True Positives are usually defined up to a small location error, as matching the ground truth perfectly would be too stringent. In our case, the spots are small compared to the object size so that the position error remains very small even in the cases where the intersection between the predicted and ground truth spot is the smallest possible one (one pixel) (see Figure 7). Therefore, we consider any overlap between a prediction and a ground truth spot to be a True Positive, as long as this ground truth spot has not been predicted before. If this is the case, the prediction is considered to be a false positive. Predicted objects that do not overlap ground truth spots are also considered to be false positives. Finally, objects in the ground truth heatmap that are not predicted by the model are considered to be false negatives. True negatives are ill-defined in object detection, and are not considered. With those three values, we can compute Precision and Recall.

**Figure 7.**
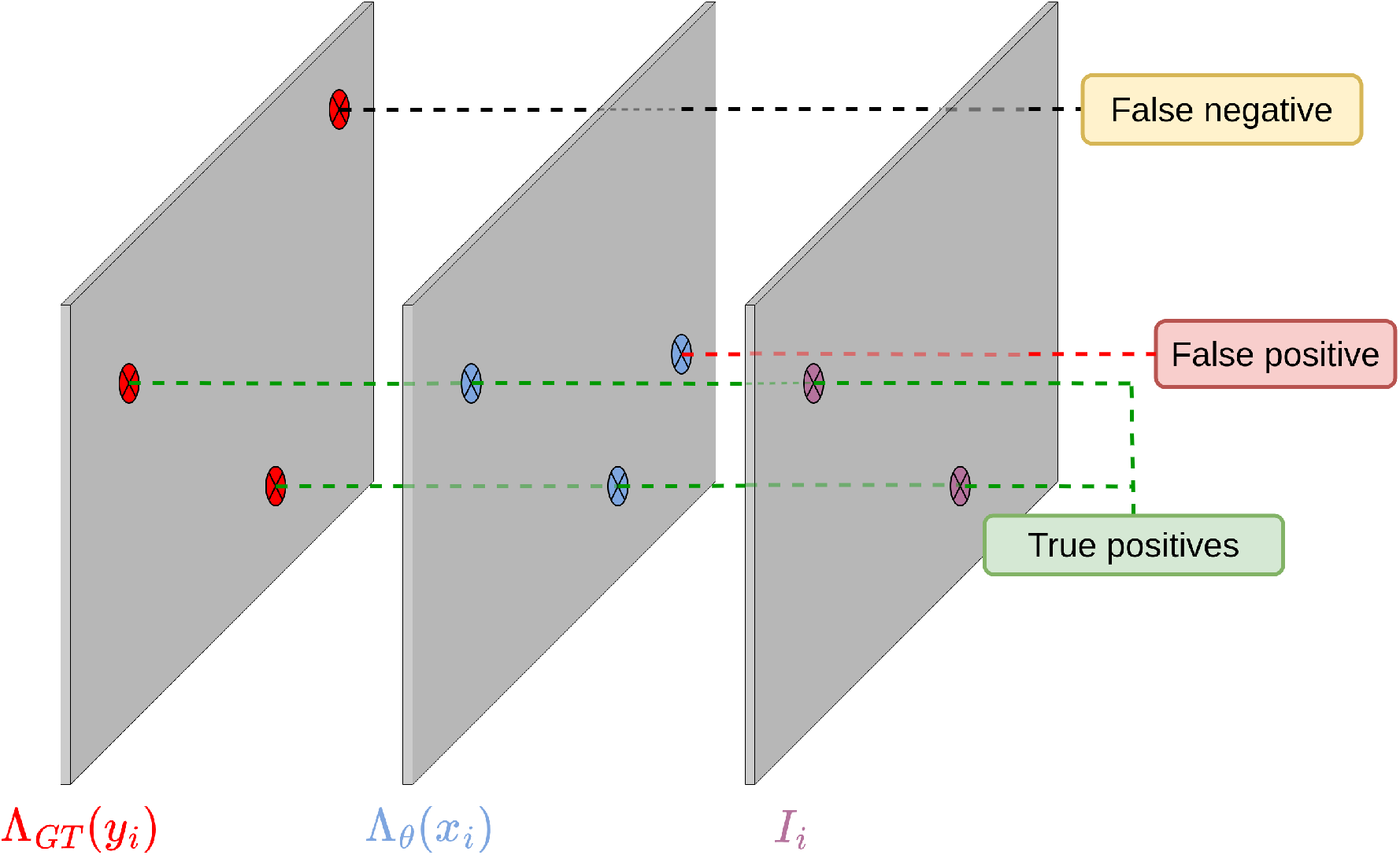
Sketch of model evaluation. The intersection ***I***_*i*_ between the binarized ground truth **Λ**_*GT*_ (*y*_*i*_) and the binarized prediction map **Λ**_*θ*_(*x*_*i*_) is computed. Objects appearing in both are True Positives, objects appearing only in **Λ**_*θ*_(*x*_*i*_) are false positives, objects appearing only in **Λ**_*GT*_ (*y*_*i*_) are false negatives. In this case, we have two True Positives, 1 False negative and 1 False positive, so that Precision is *TP/*(*TP* + *TP*) = 2*/*3 and Recall is *TP/*(*TP* + *FN*) = 2*/*3

More formally, the intersection image ***I***_*i*_ between **Λ**_*θ*_(*x*_*i*_) and **Λ**_*GT*_ (*y*_*i*_) is computed as the pixel-wise product of ground truth and prediction

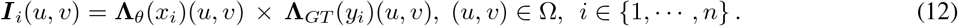

Finally, we compute the number of connected components *N*_*cc*_(***I***_*i*_), *N*_*cc*_(**Λ**_*θ*_(*x*_*i*_)) and *N*_*cc*_(**Λ**_*GT*_ (*y*_*i*_)) in ***I***_*i*_, **Λ**_*θ*_(*x*_*i*_) and **Λ**_*GT*_ (*y*_*i*_), respectively. The number of True Positives, False positives and False Negatives are defined as follows:

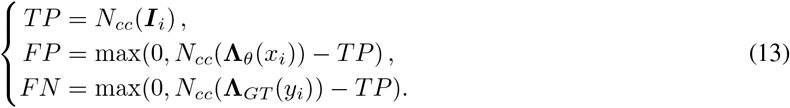

Hence, we compute Precision 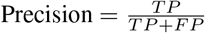 and Recall 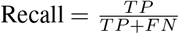 for a single (**Λ**_*θ*_(*x*_*i*_), *y*_*i*_) pair. Note that Precision and Recall are functions of the chosen confidence level *T*_*C*_; a higher confidence threshold increases Precision but decreases Recall. In what follows, we will then report results for a set of confidence thresholds 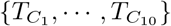 to analyze this tradeoff. Moreover, we decided to report the Precision and Recall scores separately instead of providing an aggregated metric like Average Precision (AP), as the tradeoff between those metrics in the context of biological dosimetry is especially important.

Because training is a stochastic process, the maximum predicted probability over the test set is not exactly 1; different models may get a different maximum confidence value. In other words, two sets of weights *θ* and *θ*^*′*^ will give two different maximum probabilities *p*_*θ*_ and *p*_*θ*_^*′*^, so that for each model, the performance metrics are computed over different confidence thresholds. Therefore, each performance metric curve is linearly interpolated over a common confidence grid {0.1, 0.2, *· · ·*, 0.9}. For the set of confidence thresholds that exceed the maximum probability over the complete test set, Precision is not defined. In this case, it is arbitrarily chosen to be 1 (and Recall is 0). To provide a metric showcasing performance variation across training, we reported performance quantiles (5% and 95%) for each threshold. This means that, for each threshold, 50 Precision and Recall values are computed (one for every considered epoch) and the aforementioned quantiles of those values are reported.

#### 4.2.2 Performance of model ensemble

For the ensemble, we use the same evaluation procedure as in the single model case (described in Section 4.2.1), but with the agregated binary decision map ***D***_*i*_ described in Section 3.2. However, as explained in Section 3.2, the performance of the ensemble depends on an agreement threshold *T*_*A*_ and a confidence threshold *T*_*C*_. Therefore, instead of a Precision curve, we get a Precision surface, which makes comparison with the single model case more difficult. Instead, we set a specific vote threshold, and report the same Precision curve as in the single model case. Finally, to evaluate the sensitivity of the performance to the sampling of the ensemble, we sampled 100 random ensembles, and computed *q*_05_ and *q*_95_ for every confidence threshold in the grid mentioned in the Section 4.2.1.

## 5 Experimental results

In this section, we discuss model performance, both in term of object detection and calibration curve estimation. First, we discuss the results of the single-model training and the impact of regularization term. We also show a visualization which suggests that our model is robust to the presence of debris, without the need for specific labelling or architectural choices. Finally, we discuss the performance improvements obtained with the ensemble approach, and we provide PCA-based approach to visualize model feature trajectories during training.

### 5.1 Performance of single model

In Table 2, the single non-regularized model already achieves significant gain in terms of Precision and Recall when compared to Metafer in the case of monocentric versus dicentric classification (Metafer is not designed to detect fragments), although some performance variation is noticeable during training, as confirmed by the inter-quantile range of performance. This variation suggests that there is sufficient parameter space exploration to get enough prediction diversity for aggregation to be worthwhile.

**Table 1:** Comparison between model 1 (*T*_*C*_ = 0.6, *λ* = 0), model 2 (*T*_*C*_ = 0.6, *λ* = 0.2, *ρ* = 0.1) and ensemble (4 votes, *T*_*C*_ = 0.5) for the fragment class. [*q*_5%_, *q*_95%_] interval is reported in parenthesis. For model 1 and model 2, this interval is computed over the last 50 epochs of training. For the ensemble, it is computed over a 100 randomly sampled ensembles (ensembles are sampled from checkpoints during training). All performance metrics are computed on the test set.

| | model 1 (unregularized) | model 2 (regularized) | ensemble (3 votes, $T_C = 0.5$) |
| --- | --- | --- | --- |
| Precision (frags) | 70.8% (62.0, 77.4) | 79.3% (71.8, 84.2) | 81.0% (78.5, 83.3) |
| Recall (frags) | 49.4% (37.8, 59.7) | 55.0 % (46.0, 66.3) | 65.5 % (62.4, 68.5) |

**Table 2:** Comparison between model 1 (*T*_*C*_ = 0.6, *λ* = 0), model 2 (*T*_*c*_ = 0.6, *λ* = 0.2, *ρ* = 0.1), **ensemble** (4 votes, *T*_*C*_ = 0.5), and Metafer for the dicentric class (where Metafer performance is available). All performance metrics are computed on a separate test set. Metafer performance is retrieved from previous work, and was not evaluated on the dataset used in this paper. Metafer performance should only be taken as a rough point of reference, see Section 5.1 for additional discussion of this point.

| | model 1 (unregularized) | model 2 (regularized) | ensemble (3 votes, $T_C = 0.5$) | Metafer |
| --- | --- | --- | --- | --- |
| Precision (dics) | 76.6% (68.8, 84.7) | 83.6% (75.4, 92.2) | **85.8%** (83.7, 88.3) | $\sim 40\%$ |
| Recall (dics) | 45.2% (31.6, 56.4) | 51.2% (37.0, 62.7) | **61.7%** (57.1, 65.8) | $\sim 35\%$ |

Metafer relies on conventional computer vision techniques. The chromosome objects in the metaphase are first segmented and then classified. Segmentation errors can lead to classification errors, for example when one chromosome is split during segmentation. Overall, the performance of both tasks (segmentation and classification) is not very robust to variations in image quality induced by variations in acquisition circumstances like illumation or staining quality. For example, in the segmentation task, the Recall of Metafer lies between 35% and 75%. For the classification task, The Recall is around 35%, and the Precision around 40%. Note that the performance figures given for Metafer in Table 2 are not computed on the dataset mentioned in Section 4.1, but taken from [41] instead. Although the performance of Metafer is uncertain, and we did not compute it on our dataset, we feel confident that our model brings a very significant improvement in chromosomal aberration, as the uncertainty around the ensemble performance is low enough that even in the worst case scenario, it achieves very significant performance improvements over Metafer.

Finally, training with a loss that promotes sparsity improves Precision and Recall for fragments and dicentrics, as seen in Figure 8. this improvement in precision is especially relevant for automated use, as keeping the number of false positives low is required to avoid overloading care facilities. Because all chromosomes may not be correctly retrieved (Recall is less than 1), this model tends to underestimate doses.

**Figure 8.**
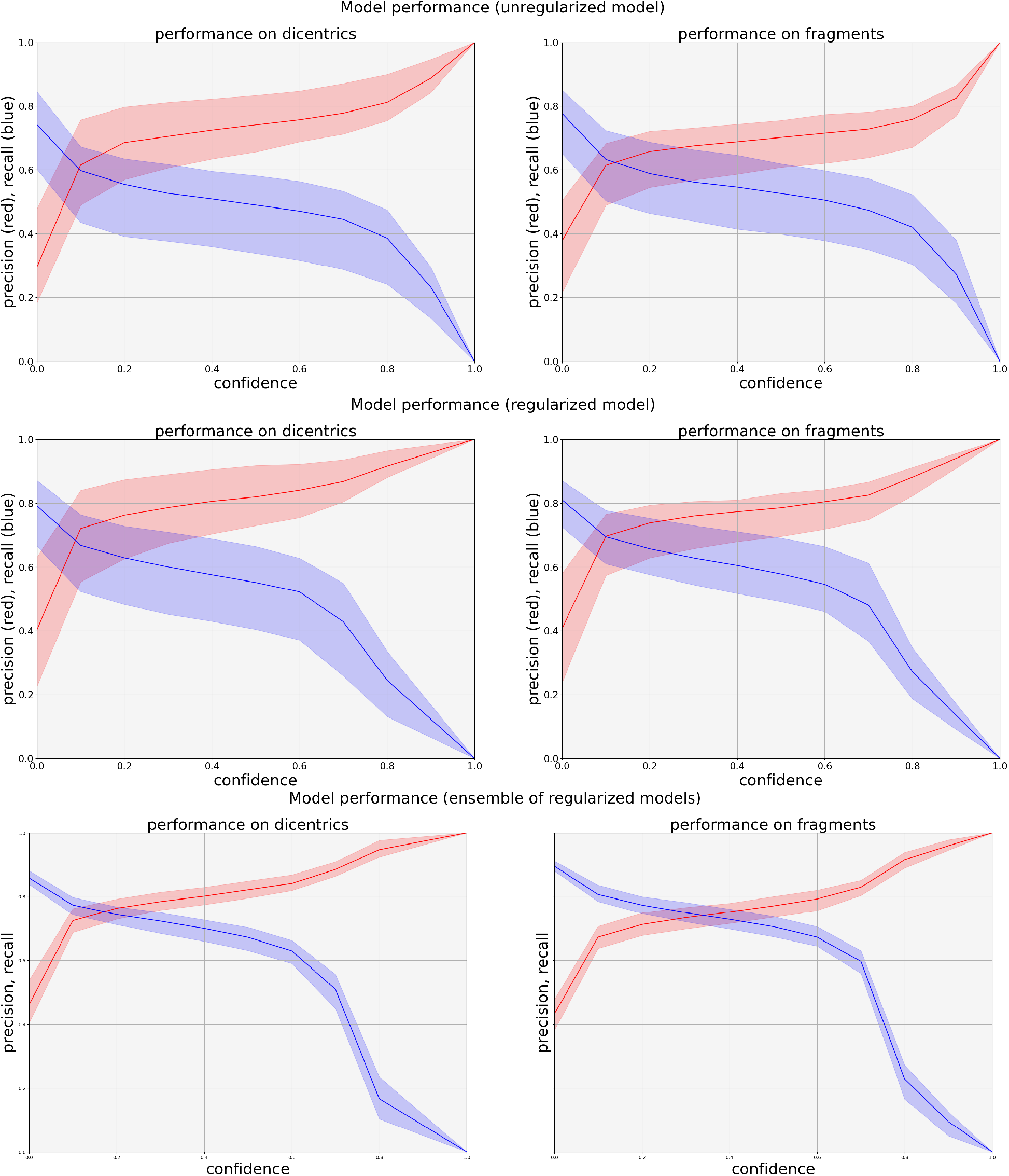
Precision, recall and False Discovery Rate (FDR) as functions of confidence for dicentrics (left) and fragments (right). Top: performance summary for the unregularized model (i.e *λ* = 0 for the sparse variation term). Middle: performance summary for *λ* = 0.2, *ρ* = 0.1. Bottom: performance summary for the ensemble of checkpoints from the training of the regularized model for a threshold of 2 votes. Shaded area indicates the [*q*_0.05_, *q*_0.95_] inter-quantile interval, computed respectively over the last 50 checkpoints for single models, and over a 100 random samples of 10 checkpoints for the bottom plot (ensemble).

### 5.2 Performance of model ensemble

In this section, we report the results obtained with the ensemble procedure. The parameter space of the model is explored through SGD. Although the selected checkpoints reach a similar validation performance, the predictions they yield vary, as shown by Figure 9. As mentioned earlier, this can be used to filter spurious predictions, as those are less likely to be present in all models of the ensemble.

**Figure 9.**
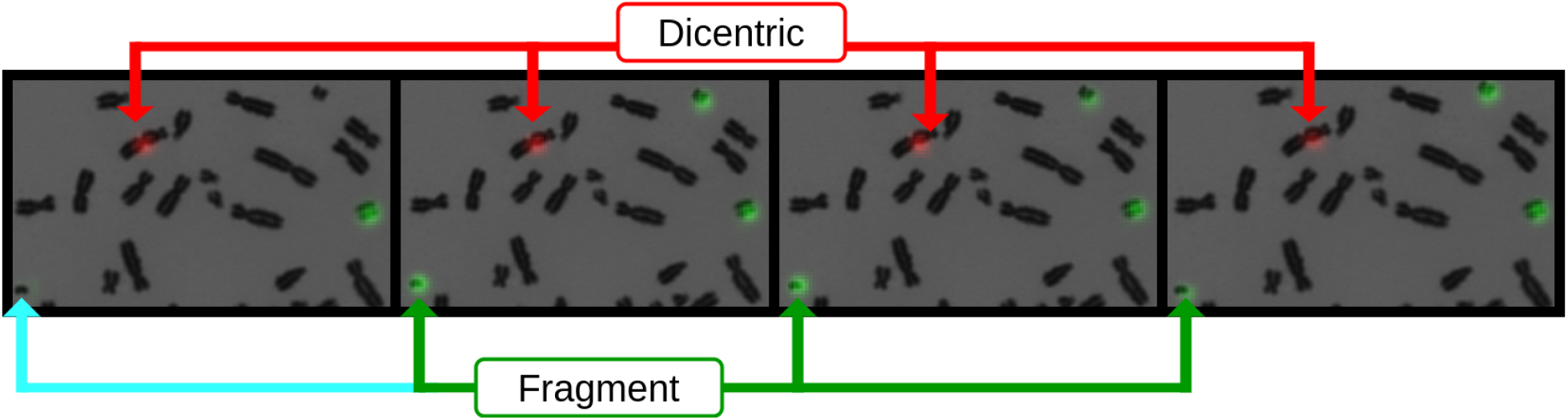
Prediction diversity for a set of four different models. The first model does not predict the fragment in the bottom left corner (a very low confidence threshold would be required to consider it as a detection), but the three next model do. All four models predict the dicentric chromosome near the center of the image correctly.

Precision is a monotonously increasing function of voting and confidence thresholds, while Recall decreases with higher confidence and higher voting thresholds. End-users may fine-tune the balance between both of those metrics depending on their goal by choosing a specific (*T*_*C*_, *T*_*A*_) pair. To estimate the sensitivity of Precision and Recall to the sampling of the checkpoints, we evaluate those metrics for 100 samples ensembles and report the *q*_05_, *q*_95_ interval for Precision and Recall in Figure 8).

To ensure that ensemble results are easily readable, we do not report surfaces for Precision and Recall. Instead, we select a single voting threshold and report the results over all agreement thresholds, like with the single model results. The ensemble provides a significant performance improvement over the single-model baseline; aggregation does help to filter out spurious detections and improves Precision and Recall, as reported in Tables 1 and 2. We chose the (*T*_*C*_, *T*_*A*_) parameters to keep Precision broadly similar across all models, for dicentrics and fragments. This makes Recall improvements more salient, but one could choose other values for confidence and vote thresholds. Overall, there is a wide set of threshold combinations that yield large performance improvements over the Metafer baseline. Furthermore, ensembling also reduces performance variation: the performance is closer between different ensembles than between single checkpoints. This suggests that our results are not dependent on a specific sampling or selection of the ensemble.

### 5.3 Robustness to non-chromosome objects in metaphase images

An automated dosimetry system should always distinguish between chromosome and non-chromosome (nuclei, debris) objects. Current automated methods reject non-chromosome objects using explicit shape analysis. For example, a nuclei can be rejected using the fact that it is broadly circular and has a uniform texture. However, the shape of debris is usually more complex, which makes metaphase selection more difficult in most dosimetry systems.

In our approach, we do not detect debris and nucleis explicitely, as they are not annotated in our dataset. Instead, Unet learns to reject debris from the training data, without the need for specific annotations or handcrafted object detection algorithms. In the rest of this section, we propose a simple visualization of this fact, using PCA.

For an image *x*_*i*_ and a set of parameters *θ*, the activation volume provided by the *ℓ*-th layer of the neural network is defined as:

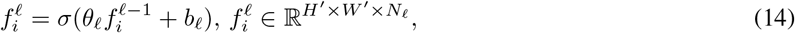

where *N*_*ℓ*_ the number of convolution filters in layer *ℓ*, i.e the number of channels of the output of this convolution layer. For a set of *n* images {*x*_1_, *· · ·, x*_*n*_}, the corresponding activation volume is of size ℝ^*n×H′ ×W′ ×N*_*ℓ*_^ This volume can be flattened in a matrix 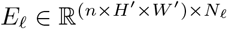, containing *n × H*^*′*^ *× W*^*′*^ samples (rows) of a random vector of size *N*_*ℓ*_. Once *E*_*ℓ*_ is centered and standardized, the eigenvectors of 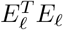 form the usual PCA orthogonal basis 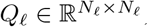, so that 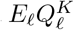 (with 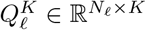 projects each pixel of the activation volume (*N*_*ℓ*_-sized vectors) onto the first *K* components (columns) of this basis.

By choosing *K* = 3 and normalizing the projected vectors to sum to one, we can plot a visualization of the activations in RGB space (Figure 10). Note that a different *Q*_*ℓ*_ is computed for each layer considered in Figure 10. Therefore, the colors do not have any specific meaning. The second image of Figure 10 from the left shows that the projection of feature vectors (retrieved at the first encoder layer) belonging to chromosomes and nuclei on the PC basis are highly similar, as the color of those regions is identical. The main reason is that the convolution filters in early layers have a small receptive field (see [2]) and mostly capture texture and edge information, which is close for subsets of chromosomes and nuclei. In the early stages of the encoder, non-chromosome objects are not differentiated from chromosomal objects.

**Figure 10.**
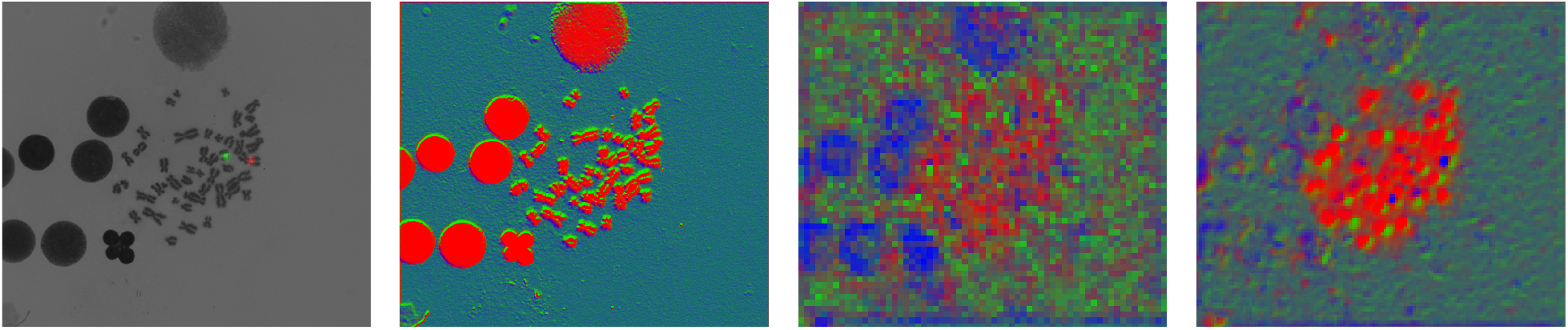
Rejection of nuclei depending on model layer. First image from the left shows the input image and ground truth Gaussian heatmap. Second image shows PCA embedding of features at the output of the first encoder layer. Third image shows PCA embeddings of features at the last encoder layer. Rightmost image shows embeddings of features at the first decoder layer. The embedded feature maps are resized from *H*^*′*^, *W*^*′*^ to *H, W* so that every image has the same size.

The third and last images of Figure 10 from the left show that in the deepest layer of the encoder, the feature vectors belonging to nuclei and chromosomes are mapped to different principal components, corresponding respectively to blue and red pixels. By stacking convolutions the model aggregates information from a larger subset of the image (the receptive field increases), which is suitable to distinguish chromosomes from nuclei on the basis of their differences in shape. This is a first indication that our model learned to reject debris and nuclei without domain knowledge, which is further confirmed by our performance results.

### 5.4 Visualization of training trajectories

In this section, we discuss the visualizations produced by the method described in Section 3.4. In Figure 11, we see the latent space of Unet at 6 different epochs. Red points on the scatterplot represent locations containing dicentric chromosomes, blue points fragments and gray points are background locations. Because those snapshot are taken at regular intervals at the end of training, classes are well separated. We see that the decision boundary of the SVM classifier changes from timestep to timestep. Some locations remain well separated in latent space from others, but this is not true for all samples. Regions in the latent space where classes overlap correspond to areas of uncertainty.

**Figure 11.**
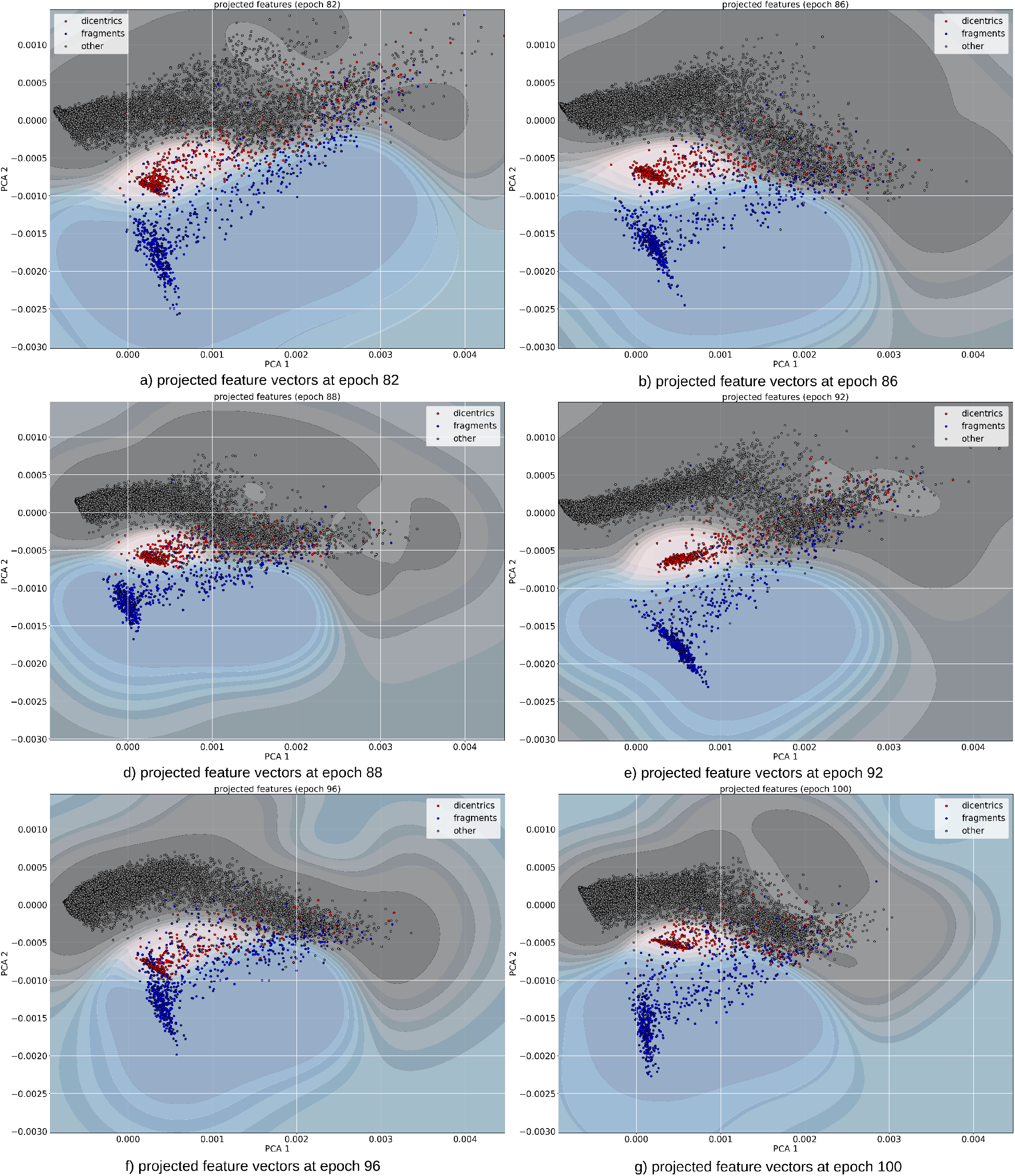
Snapshot of the training trajectory in feature space. Each point of every scatterplot represents a fixed location in an image (dicentric, fragment or background). Because of the stochasticity of training, the corresponding feature vector moves in feature space. By aligning the feature sets produced by the model at different epochs and computing a PCA dimension reduction, we can visualize the displacement of those feature vectors in feature space during training. The contour map showcases the decision boundary of a kernel SVM classifier trained to predict the type of feature vector depending on its location in feature space. While some regions of feature space remain ambiguous during training, classes tend to stay clustered together during training.

While locations with high prediction uncertainty are hard to detect because neural networks tend to be overconfident [13], an ensemble of models can be used to retrieve this information. This fact is also visible in the latent space of Unet, as shown by Figure 12. In this Figure, we performed K-Means classification on our bag of features in the latent space for the last considered epoch in Figure 11. Once this is done, we plot the trajectories of the barycenters of those clusters over training epochs. As we see, most cluster centers are well separated across epochs. This is reflected by the fact that if we consider the average SVM decision boundary across training epochs (as described in Section 3.4), most feature vectors are in low-entropy regions. Feature vectors belonging to high-entropy regions correspond to uncertain detections can be filtered as described in Section 3.2.

**Figure 12.**
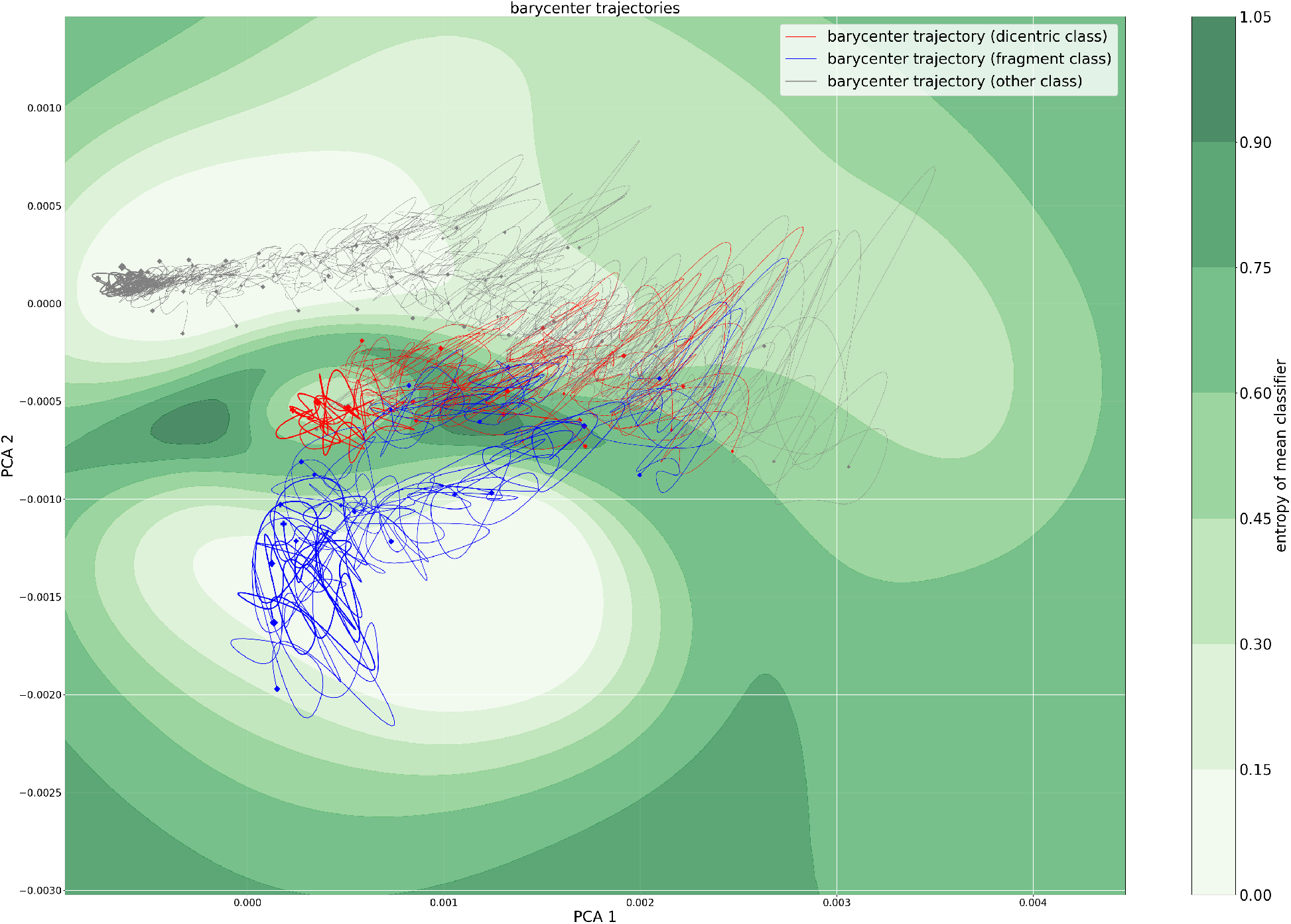
Training trajectories of feature barycenters. The scatterplots displayed in Figure 11 are clustered with *k-means* to simplify visualization. The trajectories of barycenters during training are displayed in this figure. The thickness of the trajectory shows the number of feature points in the barycenter.

### 5.5 Transfer between training and calibration curve datasets

As explained in Section 1, most images routinely analyzed by biologists do not contain any aberration, even for high doses. Our annotated training dataset is a subset of a much larger archive of patient data. This annotated dataset contains 5,430 images depicting at least one aberration over *∽* 80k images. This reduces training time considerably, and prevents the discovery of trivial models where no object is ever predicted. However, this also means that our training dataset is not an accurate representation of real-world metaphase images as metaphases containing aberrations are considerably over-represented. In the previous section, we saw that the model described in this paper performs very well in terms of object detection metrics. However, we still need to investigate wether a model trained on this unbalanced dataset can accurately estimate aberration counts on a realistic dataset.

Initial calibration curve estimates were unsatisfying: our ensemble would overestimate low doses, and underestimate high doses. Two additional details were needed to improve performance. First, we investigated the Cumulative Distribution Function (CDF) of the maximum probabilities predicted by the members of the ensemble for the dicentric and fragment class. Those CDFs were estimated using the 4 Gy subset of the calibration curve dataset, and are shown in Figure 13. Instead of setting a single threshold for all model, we picked a quantile, and retrieved the corresponding CDF value for each model. Furthermore, we used domain knowledge to reject spurious dicentric detection. We considered dicentric detections if and only if at least one fragment was present in the same image. As usual in biological dosimetry, we fitted a linear-quadratic model to the point cloud of average dicentric count retrieved on the calibration curve dataset. This led to a large improvement in our calibration curve estimation, as shown in Figure 14. Furthermore, the Metafer calibration curve is semi-automated: dicentric chromosomes undergo a manual review, because of the very high FPR of the algorithm, as described in Section 5.1. On the contrary, our calibration curve is obtained in a fully automated setting.

**Figure 13.**
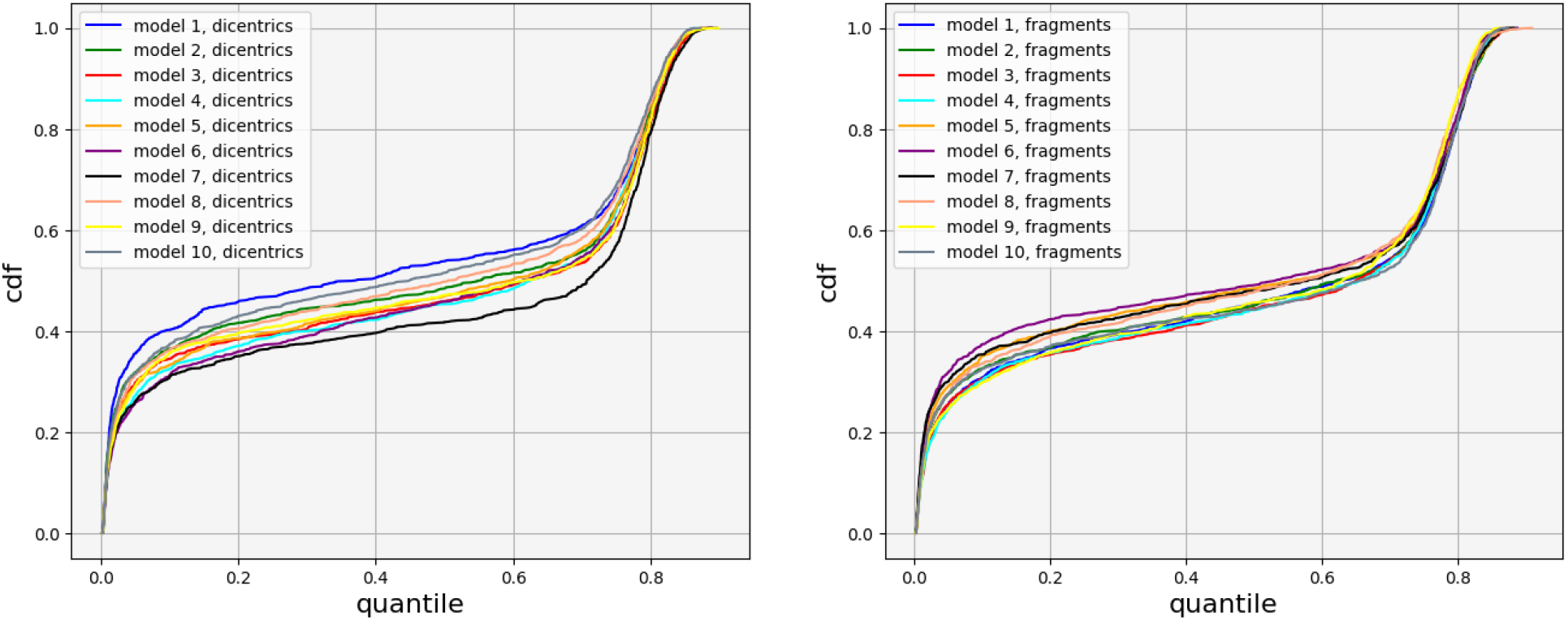
Cumulative Distribution Functions (CDF) of the maximum probabilities predicted by every member of the ensemble for the dicentric (left) and fragment (right) class over all images corresponding to a 4 Gy dose in the calibration curve dataset.

**Figure 14.**
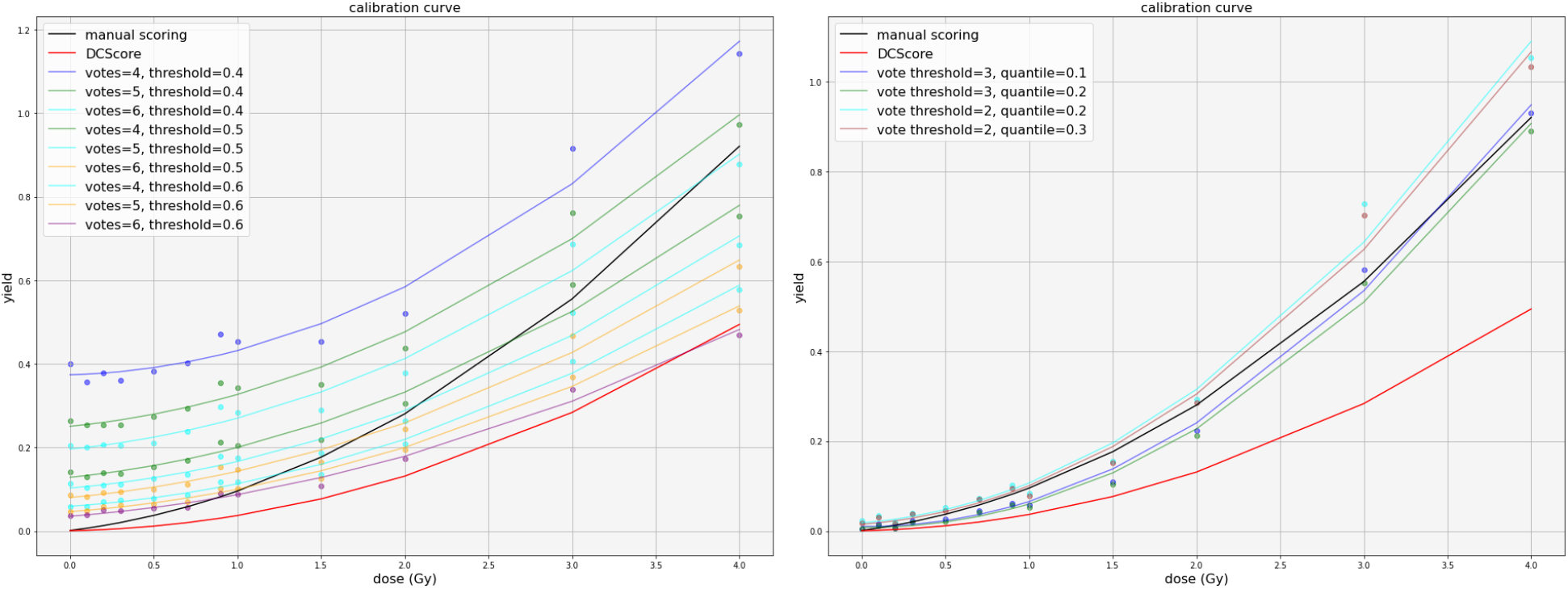
Calibration curves estimated by the ensemble. Left: calibration curve before setting a threshold per model and using domain knowledge. Right: calibration curve after model-adaptive thresholding and using domain knowledge. To improve readability, we show the 4 curves closest to the manual calibration curve displayed in black. Metafer curve is displayed in red.

## 6 Discussion and future works

In biological dosimetry, estimating the average number of chromosomal aberrations per peripheral blood lymphocyte is necessary to estimate an ionizing radiation dose. However, human expertise is required and is therefore a bottleneck to scale chromosome counting beyond a few hundred images per patient.

In this paper, we evaluated Unet as an aberration detection model for biological dosimetry. Unet outperformed Metafer (a current commercial solution) in terms of Precision and Recall by a wide margin. Our approach is learning-based and differs significantly from the current state of the art in terms of how much domain knowledge of chromosome morphology is incorporated. We demonstrated a high level of performance without the need for significant shape modeling. Feature visualization suggests that the model learns to reject debris and nuclei in an unsupervised manner, without the need for object-specific annotations for monocentric chromosomes or debris. Furthermore, a simple regularization term modeling the intrinsic heatmap sparsity helps performance.

We pushed this performance further by ensembling several checkpoints collected during training. We proposed a visualization of the latent features of Unet during training to explore the relationship between the dynamics of training and this performance improvement. Furthermore, we showed that the variation in performance between different (randomly sampled) ensembles is lower than between single checkpoints of a training run. This is especially relevant in the context of the deployment of a deep learning model in an automated fashion in a medical setting. Those improvements can be achieved without the need for any architectural modifications or extensive hyperparameter calibration.

Our database is very imbalanced: the average number of aberration per cell is over 1, which corresponds to an extremely high dose of ionizing radiation. It is very likely that some of the spurious detections can be attributed to this training set imbalance. We evaluated our ensemble of Unet in a realistic setting, on a calibration curve dataset. Using model-adaptive thresholding and the domain knowledge of co-occurence of dicentrics and fragments, we reach a very competitive calibration curve, widely outperforming the Metafer baseline. Furthermore, our ensemble can be used in a fully automated fashion, while the Metafer solution requires manual review to reduce the number of false positives. It is therefore possible to build a competitive aberration detection system even with a large distribution gap between training and inference images.

## Data and software availibity

The data cannot be made publicly available due to restricted access under IRSN ethics and security policy, and because informed consent from participants did not cover the publication of this data.

## Acknowledgements

This work was jointly supported by the Defense Innovation Agency (AID) and the National Research Agency (Increased ANR-20-ASTR-0005, France-BioImaging ANR-10-INBS-04-07). Additionally this research was funded in part by IRSN and Région Bretagne.

## Author contributions

M.A.B and C.K. devised the project and the main conceptual ideas, supervised the project and was in charge of overall direction and planning. A.D. designed and implemented the method, in discussion with E.G, J.S.M., A.V., P.F. D.F., L.B. M.V., and G.G. who designed and provided the datasets. A.D. implemented the analysis workflows applied to real images, in discussion with E.M. A.D, C.K., A.M.B., and E.M. co-wrote the manuscript. All authors provided critical feedback and helped shape the research, analysis and manuscript

